# Mechanistic insight into the ATP hydrolysis cycle of tick-borne encephalitis virus helicase

**DOI:** 10.1101/2022.03.15.484399

**Authors:** Paulina Duhita Anindita, Marco Halbeisen, David Řeha, Roman Tůma, Zdeněk Franta

**Author notes:** Corresponding author: Zdeněk Franta.

## Abstract

Helicase domain of nonstructural protein 3 (NS3H) unwinds double-stranded RNA replication intermediate during flavivirus life cycle in ATP-dependent manner. While the helicases mechanism of Dengue and Zika viruses has been extensively studied, little is known about helicase activity of the tick-borne encephalitis virus NS3. In the current study, we demonstrated that ATP hydrolysis cycle of NS3H is strongly stimulated by ssRNA but not ssDNA, suggesting that NS3H is an RNA-specific helicase. However, ssDNA binding inhibits ATPase activity in a non-competitive manner. We also captured several structural snapshots of key ATP hydrolysis stages. An intermediate, in which the inorganic phosphate (Pi) and ADP resulted from ATP hydrolysis and remain trapped inside the ATPase site, suggests that Pi release is the rate-limiting step and is accelerated by RNA binding and/or translocation. Based on these structures, we modeled NS3H-ssRNA and -ssDNA binding and performed MD simulations. Our model suggests that NS3H-ssRNA binding triggers conformational changes, revealing the coupling between helicase and ATPase activities. The structural models revealed that ssDNA inhibition may occur via non-specific ssDNA binding to several positively charged surface patches which in turn causes repositioning of ATP molecule within the ATPase site.

## 1. Introduction

Tick-borne encephalitis virus (TBEV) is an enveloped single-stranded (ss), positive-sense RNA virus from the family *Flaviviridae*, genus *Flavivirus*. The virus is neurotropic and causes tick-borne encephalitis which affects mostly adult population within European and north-eastern Asian countries (1). While TBEV vaccines are available due to the lack of targeted campaigns the actual vaccination coverage is low even in areas with high risk. There is currently no specific treatment available (2,3). To develop antivirals against TBEV, it is important to understand the structure and function of viral enzymes that are involved in the replication.

TBEV encodes three structural and seven non-structural proteins (NS). Among the NS proteins, NS3 plays a role as a multifunctional protein which comprises two domains: protease and helicase domain. Protease domain is located at the N-terminal NS3 (172 aa). It is a chymotrypsin-like serine protease responsible for viral polyprotein processing. This domain is connected by a short flexible linker to the C-terminal helicase domain (NS3H, 434 aa) which exhibits several activities: nucleoside 5’-triphosphatase (NTPase), RNA helicase and RNA 5’-triphosphatase (RTPase). The RNA helicase activity uses energy from NTP hydrolysis for unwinding of dsRNA replication intermediates while RTPase removes the terminal γ-phosphate from the 5’-triphosphate end of the positive-sense ssRNA before mRNA capping by the methyl transferase domain of NS5 protein (4,5).

In this study, we focus on NS3H which is a monomeric enzyme belonging to the DEAD/H box sub-family within the helicase superfamily 2 (SF2) (6,7) and consisting of three subdomains. Subdomain 1 and 2 both exhibit the highly conserved Rec-A like fold typical of P-loop NTPases. The NTPase active site is located in a cleft between the subdomains 1 and 2, and involves motifs I (Walker A), II (Walker B) and VI. A groove between subdomain 3 and subdomain 1 and 2, respectively, forms the RNA binding site. Helicase activity relies on coupling between NTPase and RNA binding sites (8,9).

Several structural and functional studies on flavivirus helicases from DENV, ZIKV, YFV, Kunjin virus and JEV have been previously reported and revealed high structural conservation among them (10-14). Previous biochemical and structural studies have provided insight into substrate binding and indicated structural changes and RNA interacting residues associated with ATP hydrolysis cycle of DENV helicase (11,15). Recent molecular dynamics (MD) simulations (9,16) provided further insight into conformational changes associated with RNA binding and stimulation of the ATP hydrolysis which was considered as the rate-limiting step. In contrast, phosphate (Pi) release has been established as the rate-limiting step for HCV NS3 helicase that is considered a mechanistic model system for SF2 superfamily (17). Hence, it remains to be seen whether Pi release or the hydrolysis is the rate-limiting step and whether the coupling and the associated allosteric changes are conserved among all SF2 helicases and in particular between helicases from different flaviviruses.

Here we biochemically characterized helicase from TBEV and obtained structures of key intermediates along the ATPase cycle (nucleotide-free, apo; ADP; adenylyl-imidodiphosphate, AMPPNP; and hydrolysis product, ADP-Pi). While the overall structure of the TBEV helicase is closely related to that of DENV and ZIKV structural variation is observed in different nucleotide states. Trapping of ATP hydrolysis products within the crystal and the negligible basal activity (i.e., in the absence of RNA) suggest that the ATPase rate limiting step is phosphate release which is allosterically stimulated by RNA. Indeed, ssRNA binds to NS3H with nanomolar affinity and stimulates its ATPase activity. We further observed that ssDNA inhibits the RNA-stimulated ATPase activity and explored ssRNA and ssDNA binding by modeling and MD simulations. The models and binding assays suggest that ssDNA may exert inhibition not by direct competition with ssRNA but by binding to several positively charged patches on the surface. Such DNA binding leads to destabilization of ATP binding and repositioning in the binding cleft, and indirectly interferes with RNA binding.

## 2. Experimental procedures

### a. Plasmid construction and protein production

A DNA fragment encoding the full-length helicase domain including the linker region between protease and helicase domains (amino acid residues 173-621) from TBEV strain HYPR, hereafter referred as NS3H, was amplified from pUC57-HYPR-FRAGII (kindly provided by Prof. Daniel Růžek, State Veterinary Institute, Czech Republic) using Q5 polymerase and gene specific primers (Table S1). The PCR product was cloned into pET-19b plasmid (Novagen) and sequence verified. Recombinant NS3H was produced in *E. coli* BL21-CodonPlus (DE3)-RIPL competent cells (Agilent Technologies). Briefly, transformed competent cells were grown at 37°C in LB media supplemented with 100 μg/ml ampicillin and 35 μg/ml chloramphenicol until OD_600_ reached 0.5. The cells were chilled for 30 min at 4°C and the protein production was induced with 1 mM IPTG followed by incubation for 20 h at 18°C. The cells were harvested by centrifugation at 4,000 *g* for 30 min at 4°C and stored at -80°C prior further use.

### b. Protein purification

Cells were resuspended in buffer A (0.02 M sodium HEPES pH 7.0, 0.5 M NaCl, 10 μg/ml DNAse I) and lysed using LM20 Microfluidizer® Processor (Microfluidics). Resulting cell lysate was clarified by ultracentrifugation at 40,000 *g* for 60 min at 4°C and collected supernatant was loaded on HisTrap FF column (Cytiva) equilibrated in buffer A. Histidine-tagged NS3H was eluted using 70% of buffer B (0.02 M sodium HEPES pH 7.0, 0.5 M NaCl, 1 M imidazole). Fractions containing recombinant NS3H were pooled together, and the elution buffer was exchanged for buffer C (0.02 M sodium HEPES pH 7.0, 0.15 M NaCl) using Amicon Ultra-15 spin columns with 30 kDA cutoff (Merck). To remove the residual nucleic acids bound to NS3H, the sample was loaded into HiTrap Heparin HP column (Cytiva) pre-equilibrated in buffer C and NS3H protein was eluted using 1M NaCl. Finally, eluted NS3H was concentrated using Amicon Ultra-15 with 30 kDA cutoff (Merck) spin columns and loaded into a Superdex 200 Increase 10/300 GL (Cytiva) pre-equilibrated in buffer C for final polishing. The purity of the protein was analyzed using an SDS-PAGE gel and MALDI-TOF mass spectrometry.

### c. Crystallization, data collection and data processing

The apoNS3H crystal was obtained using a sitting drop vapor diffusion method in 96-well plates assisted by Oryx-6 crystallization robot (Douglas Instruments) at 18°C. The protein in buffer C (2.2 mg/ml) was crystallized using Ligand-Friendly Screening HT-96 (Molecular Dimensions) for the initial screening and protein crystals grew in 0.1 M Bis-Tris Propane pH 6.5, 0.2 M sodium acetate trihydrate, 20% w/v PEG 3350, 10% v/v ethylene glycol. Crystals for NS3H-ADP-Mn^2+^ complex were obtained by co-crystallization of NS3H (2.2 mg/ml) with 5 mM MnCl_2_ and 5 mM ADP (Sigma Aldrich) in 0.1 M Bis-Tris Propane pH 6.5, 0.2 M sodium acetate trihydrate, 20% w/v PEG 3350, 10% v/v ethylene glycol. Crystals for NS3H-AMPPNP-Mn^2+^ complex were obtained by co-crystallization of NS3H (2.2 mg/ml) with 5 mM MnCl_2_ and 5 mM AMPPNP (Sigma Aldrich) in 0.1 M Bis-Tris Propane pH 6.5, 0.02 M sodium potassium phosphate pH 7.5, 20% w/v PEG 3350, 10% v/v ethylene glycol. With the same condition, NS3H was co-crystallized with 5 mM ATP (Thermo Fisher Scientific) and 5 mM MnCl_2_. For data collection, each single crystal was fished out and flash-cooled in liquid nitrogen without any additional cryoprotection. X-ray diffraction intensities were collected on BL14.1 at BESSY II electron-storage ring operated by the Helmholtz-Zentrum Berlin (18). The collected data sets were processed using *XDS* (19) with the *XDSAPP* graphical user interface (20). Data collection and processing statistics are given in Table 1.

**Table 1.** Crystallographic data collection and refinement statistics.

| Data set<br>(pdb code) | Apo<br>(7OJ4) | ADP- Mn<sup>2+</sup><br>(7BLV) | AMPPNP- Mn<sup>2+</sup><br>(7BM0) | ADP-Pi- Mn<sup>2+</sup><br>(7NXU) |
| --- | --- | --- | --- | --- |
| *Data collection* |  |  |  |  |
| Space group | *P* 4<sub>1</sub> 2<sub>1</sub> 2 | *P* 4<sub>1</sub> 2<sub>1</sub> 2 | *P* 4<sub>1</sub> 2<sub>1</sub> 2 | *P* 4<sub>1</sub> 2<sub>1</sub> 2 |
| *Unit cell parameters* |  |  |  |  |
| *a=b, c* (Å), $\alpha=\beta=\gamma$ (°) | 73.30, 196.05, 90.00 | 73.12, 196.13, 90.00 | 72.95, 196.45, 90.00 | 72.98, 196.59, 90.00 |
| Resolution range (Å)<sup>a</sup> | 49.06–1.83<br>(1.94–1.83) | 45.78–2.10<br>(2.23–2.10) | 45.71–1.90<br>(2.00–1.90) | 48.81–2.10<br>(2.22–2.10) |
| Unique reflections<sup>a</sup> | 48205 (7581) | 32056 (5054) | 43006 (6796) | 31603 (5033) |
| Multiplicity<sup>a</sup> | 24.54 (17.03) | 14.31 (13.77) | 14.25 (14.64) | 13.57 (13.42) |
| Completeness (%)<sup>a</sup> | 99.9 (99.3) | 99.9 (99.7) | 99.9 (99.7) | 98.4 (99.5) |
| *I*/σ<sup>a</sup> | 28.87 (2.22) | 24.38 (2.53) | 19.10 (2.03) | 23.02 (3.21) |
| R<sub>meas</sub> (%)<sup>a</sup> | 7.5 (126.6) | 6.7 (113.3) | 8.6 (118.1) | 9.9 (83.7) |
| Wilson B-factor (Å<sup>2</sup>) | 34.3 | 47.1 | 35.6 | 34.2 |
| *Refinement* |  |  |  |  |
| R<sub>work</sub>/R<sub>free</sub><sup>b</sup> (%) | 18.1/22.9 | 23.3/25.9 | 21.6/22.9 | 21.4/25.1 |
| Average B-factor (Å<sup>2</sup>) | 46.0 | 57.0 | 42.0 | 42.0 |
| *No. of non-H atoms* |  |  |  |  |
| Protein | 3375 | 3414 | 3389 | 3357 |
| Water | 209 | 75 | 230 | 200 |
| Metal ions | - | 1 (Mn<sup>2+</sup>) | 1 (Mn<sup>2+</sup>), 1 (Na<sup>+</sup>) | 1 (Mn<sup>2+</sup>), 1 (Na<sup>+</sup>) |
| Ligands | - | 27 (ADP) | 31 (AMPPNP) | 27 (ADP), 5 (PO<sub>4</sub>) |
| *RMSDs* |  |  |  |  |
| Bond length (Å) | 0.0146 | 0.0107 | 0.0155 | 0.00091 |
| Bond angles (°) | 1.929 | 1.619 | 1.765 | 1.540 |
| *Ramachandran* |  |  |  |  |
| Favored (%) | 97 | 95 | 97.37 | 97 |
| Allowed (%) | 3 | 5 | 2.63 | 12 |
| Outliers (%) | 0 | 0 | 0 | 0 |

<sup>a</sup>Values in parentheses are for the highest resolution shell.<sup>b</sup>R<sub>free</sub> was calculated using 5% of the data excluded from refinemen

### d. Structure determination and refinement

The apoNS3H structure was solved by a molecular replacement with the program *MOLREP* (21) from the *CCP4i2* suite (22) using ZIKV helicase (PDB: 6ADW) as a starting model. The ADP-Mn^2+^ and AMPPNP-Mn^2+^ ternary complexes were solved using the apoNS3H structure as the starting model. Where applicable, TLS (Translation/Libration/Screw) groups were initially identified using TLS Motion Determination (23) followed by restrained and TLS refinement cycles using REFMAC5 (24) and interspersed with manual model rebuilding using the *Coot* (25). The quality of the structure was analyzed using *MolProbity* (26). Refinement statistics and stereochemistry analysis are shown in Table 1. Superposition of structures and figures were prepared using PyMOL (27) and UCSF Chimera (28).

### e. ATPase assay

ATPase activity of NS3H was monitored using EnzChek™ Phosphate Assay Kit (Molecular Probes, Inc.) enabling the quantification of Pi release, which is translated as the rate of ATP hydrolysis by NS3H. The assay was done in a final volume of 200 μl in a 96-well plate according to manufacturer’s instruction. Briefly, 6.25 nM NS3H was mixed with individual assay components in a reaction buffer supplied by the kit and with either RNA (poly(A), 1.4 mM, Merck) or ssDNA_41_ (1 mM, Table S1), and incubated for 10 min at 30°C. The hydrolysis reaction was started by adding ATP (Thermo Fisher Scientific). Steady-state kinetics of Pi release from ATP hydrolysis was measured using a Synergy H1 microplate reader (BioTek Instruments) at 360 nm. For determination of kinetics constant of ATP substrate, the initial velocity of the reaction was calculated and expressed in micromolar concentration of Pi released per second interpolated from a calibration curve done using KH_2_PO_4_ standard solution. Data were fitted to the Michaelis equation using GraphPad Prism 9.0 (GraphPad Software, Inc.) as follows:

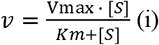

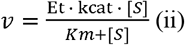

where *V*_*max*_ is the maximum velocity of enzyme turnover number, *K*_*m*_ is the Michaelis-Menten constant, *Et* is the total enzyme concentration (measured by UV absorption at 280 nm using ε _M_= 78295 M^-1^cm^-1^, estimated by ProtParams utility https://web.expasy.org/protparam/), *kcat* is the turnover number and *S* is the concentration of ATP substrate.

### f. Fluorescence anisotropy-based binding assay

Fluorescence anisotropy-based binding assay was performed by adding10 nM 6-FAM-ssRNA_12_ (Table S1) into serial dilutions of NS3H in an assay buffer (20 mM Tris-HCl pH 7.5, 100 mM NaCl, 1 mM MgCl_2_, 7.5% glycerol). The reaction mixtures (100 μl) in triplicates were incubated for 45 min at 30°C prior to measurements in ThermoScientific™ black 96-well immuno plates using Synergy H1 microplate reader. Anisotropy (r) values were calculated as follows:

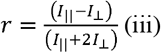

where *I*|| is the parallel emission signal and *I*. ⊥ is the perpendicular emission signal. Observed anisotropy values were plotted as a function of protein concentration. Dissociation constant (*K*_*D*_) between NS3H and RNA was determined by non-linear curve fitting of fluorescence anisotropy data to the Eq. (iv) using GraphPad Prism 9.0.

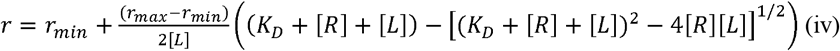

where *r, rmin* and *rmax* are the observed, the minimum and the maximum anisotropy, respectively, and [R] and [L] are the equilibrium concentration of NS3H and 6-FAM-ssRNA_12_ ligand, respectively.

Influence of DNA against NS3H-RNA interaction was examined by adding 800 nM unlabeled-ssDNA_12_, -ssDNA_20_ or -ssDNA_41_ (Table S1) into a mixture of 10 nM 6-FAM-ssRNA_12_ and 100 nM NS3H in the assay buffer.

### g. Molecular dynamics simulation

All simulations were performed using GROMACS (29) with AMBER99SB-ILDN force field (30) for proteins, nucleic acids and ions. Simulations were performed at constant temperature 300K (maintained by a modified Berendsen thermostat (31)) and pressure 1 bar (Parrinello-Rahman coupling (32)). A generalized AMBER force field (33) employing the standard procedure (34) was used for ATP. The complexes were solvated in an octagonal box using TIP3P explicit water model. Chloride and/or sodium ions and Mg^2+^ (in the presence of ATP) were added to neutralize the overall net charge. All simulations were performed for minimum of 0.5 μs or longer until stable configuration was achieved (verified by stable root mean square deviation for over 100 ns). ApoNS3H structure (this work, PDB: 7OJ4) was used and residues missing from the density maps (i.e., flexible loops) were modeled using PyMOL (22). Short (hexamer) RNA oligomers encompassing part of the conserved 5’ untranslated region (5’-UTR) from TBEV genome (5’-AGAUUU-3’) was modeled using the resolved oligonucleotide (5’-AGACUA-3’) in the crystal structure of DENV4 NS4h:RNA_12_ complex (PDB: 2JLU) (11) and protein structure superposition.

Models of heterologous non-specific surface binding were constructed using ssDNA_6_ or long ssDNA_41_ (Table S1) using initial B-form backbone and base stacking conformation. The helix was placed in vicinity at various places around the surface of RecA domains and then simulated. Simulations with the ATP placed in the binding site (using position of AMPPNP in the crystal structure PDB: 7BM0 for initial placement) were also performed to explore effects of DNA on ATP binding. Relative binding energies were computed for sub-set of the simulation frames using generalized Born surface area (GBSA) method with inclusion of the Still’s model (35) for computation of the effective Born radii implemented in the GROMACS program package (29). Superposition of structures and figures were prepared using UCSF Chimera (28).

### h. Data availability

The atomic coordinates and experimental structure factors were deposited in the Protein Data Bank under accession codes: 7OJ4 (apoNS3H), 7BLV (NS3H-ADP-Mn^2+^), 7BM0 (NS3H-AMPPNP-Mn^2+^), and 7NXU (NS3H-ADP-Pi-Mn^2+^). All remaining data are present within this article.

## 3. Results and discussion

### a. RNA-stimulated NTPase activity of recombinant NS3H is inhibited by DNA

The recombinant NS3H protein fused with N terminal 10X-histidine tag was purified to homogeneity (Fig. S1). To assess the ATPase activity of NS3H, a phosphate release assay was performed in the absence and presence of nucleic acids, ssRNA or ssDNA. ATPase activity of NS3H was strongly dependent on the presence of RNA substrate, poly(A), while the basal ATPase activity was very low (Fig. 1A). NS3H also hydrolyzed other NTP substrates (GTP, CTP, and UTP) in the presence of poly(A) with UTP being slightly less favored (Fig. 1B). This demonstrated that NS3H exhibits little specificity for different NTP substrates as shown previously for DENV helicase (36). Basic steady state kinetic parameters for ATP were determined in the presence of poly(A) (Fig. 1C), Michaelis-Menten constant *K*_*M*_ = 125 ± 15 μM, turnover *kcat* = 8.8 ± 0.2 sec^-1^, and these are similar to DENV helicase in the presence of short 5’-UTR RNA fragment (15).

**Figure 1.**
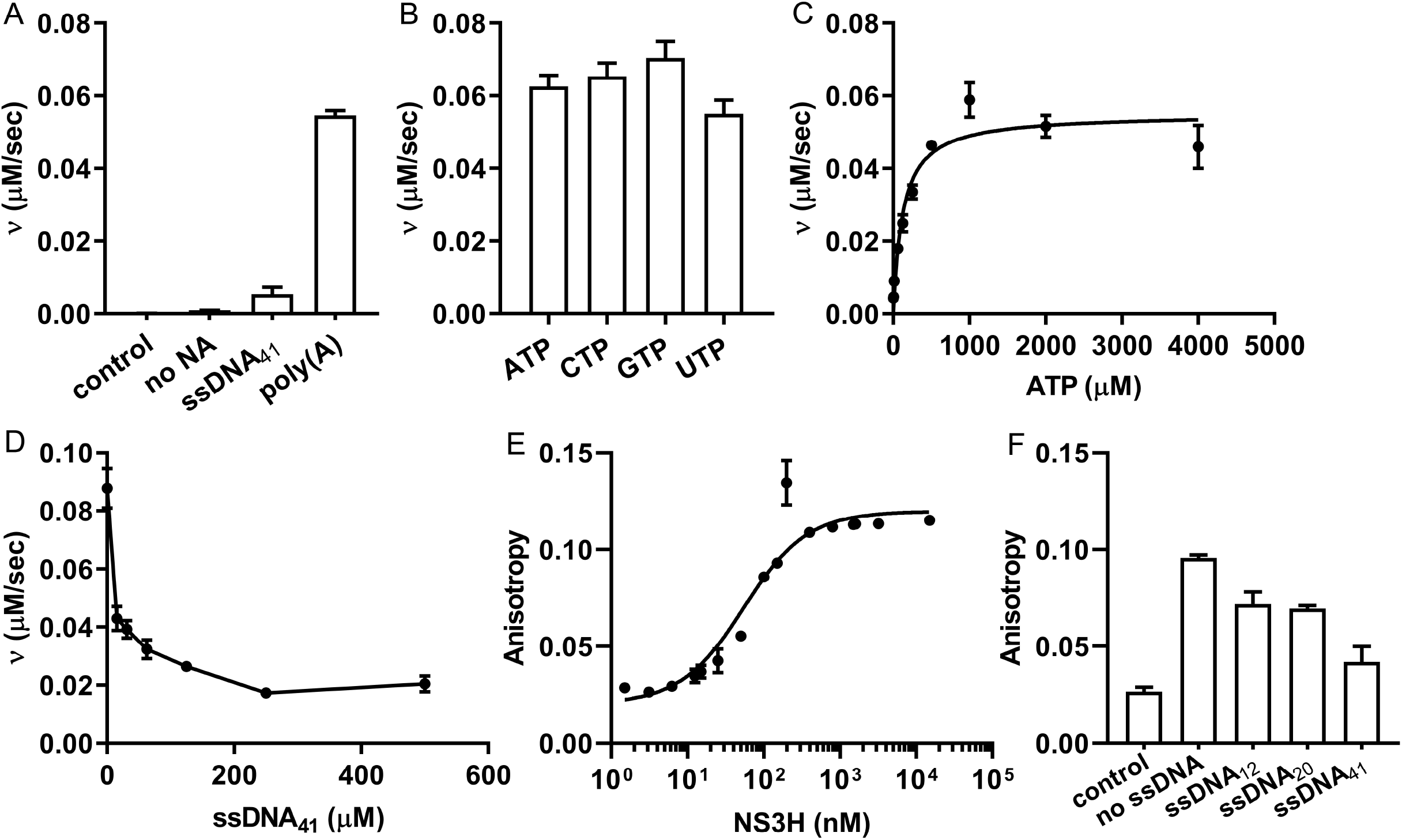
ATPase activity of recombinant NS3H. (A) ATPase activity of 6.25 nM NS3H was stimulated by 1.4 mM (nucleotide concentration) poly(A) in the presence of 1 mM ATP. The presence of 1 mM ssDNA_41_ did not show similar stimulation. The possible free Pi contamination from ATP in the absence of NS3H (control) and ATP hydrolysis by NS3H in the absence of any nucleic acid (no NA) were measured. (B) NS3H hydrolyzed 1 mM NTP substrates (ATP, CTP, GTP, and UTP) in the presence of poly(A). (C) Michaelis-Menten plot of phosphate release velocity in the presence of 6.25 nM NS3H for ATP substrate (0-4000 μM). (D) Concentration-dependent inhibition of RNA-stimulated ATPase activity by ssDNA_41_ (15.625–500 μM). (E) Binding of NS3H to 6-FAM-ssRNA_12_ measured by fluorescence anisotropy. The measurement was done using 10 nM 6-FAM-ssRNA12 and increasing concentration of NS3H from 1.5 nM to 15 μM. (F) Inhibition of 6-FAM-ssRNA_12_ binding to NS3H by ssDNA_12_, ssDNA_20_ or ssDNA_41_ was monitored using fluorescence anisotropy. The anisotropies of 6-FAM-ssRNA12 in the absence (control) and presence of NS3H (no ssDNA) were measured as controls.

While ssDNA failed to stimulate ATPase activity it can partially inhibit it in a dose-dependent manner at micromolar concentrations (Fig. 1D). However, these micromolar concentrations are much higher than affinity for ssRNA (apparent *K*_D_ = 49.7 ± 9.3 nM), which was determined by fluorescence anisotropy-based binding assay (Fig. 1E). Using the same approach, we observed that ssDNA partially competes with RNA binding at ∼20-fold molar excess of apparent *K*_*D*_ for ssRNA substrate. Longer ssDNAs inhibited RNA binding more effectively (Fig. 1F), suggesting DNA binds to several weaker binding sites. Altogether, these results suggest that the binding of the helicase to RNA substrate drives ATP hydrolysis and DNA may act as an inhibitor via non-specific binding. This specificity of NS3H toward RNA substrate is different from HCV helicase (37) and DENV helicase (38) which, in addition to RNA binding, also exhibit DNA binding with affinities ranging from nanomolar to micromolar concentration.

### b. TBEV NS3H structure is similar to that of other flavivirus helicases

The apoNS3H protein crystallizes in space group *P*4_1_2_1_2 at 1.83Å. The refined model of apoNS3H (residues 173-621) is complete except for an N-terminal 10X-histidine tag region derived from pET-19b vector together with a short flexible linker region between protease and helicase domains of NS3 (residues E173-Q181) and two non-conserved, disordered loop regions (residues P251-G259 and T502-P506). Non-protein positive *2Fo – Fc* electron density was attributed to water molecules. The resulting 3D protein structure shows overall similarity with already published helicase structures of other flaviviruses (10-14,39-41). NS3H has a clover-shaped architecture divided into three domains containing seven conserved motifs of SF2 helicase family (Fig. 2A and Fig. S2). The Rec-A-like domains 1 and 2 adopt the α/β open sheet topology (Rossman fold) (5). Domain 1 (residues W188-E329) consists of six β-strands surrounded by four α-helices, while domain 2 (residues P330-G486) consists of three α-helices and six β-strands with one anti-parallel β hairpin passing close to domain 3. Domain 3 (residues L487-R621) consists of four α-helices and one β-hairpin. To our knowledge, the apoNS3H crystal structure derived from TBEV MucAr-HB-171/11 virus strain has also been reported elsewhere (7); however, the coordinates have not been deposited in Protein Data Bank. Similar crystal structure of apoNS3H from TBEV HYPR strain is also available in the Protein Data Bank (PDB: 7JNO) with the overall RMSD of 0.8Å against our apoNS3H calculated using DALI server (42).

**Figure 2.**
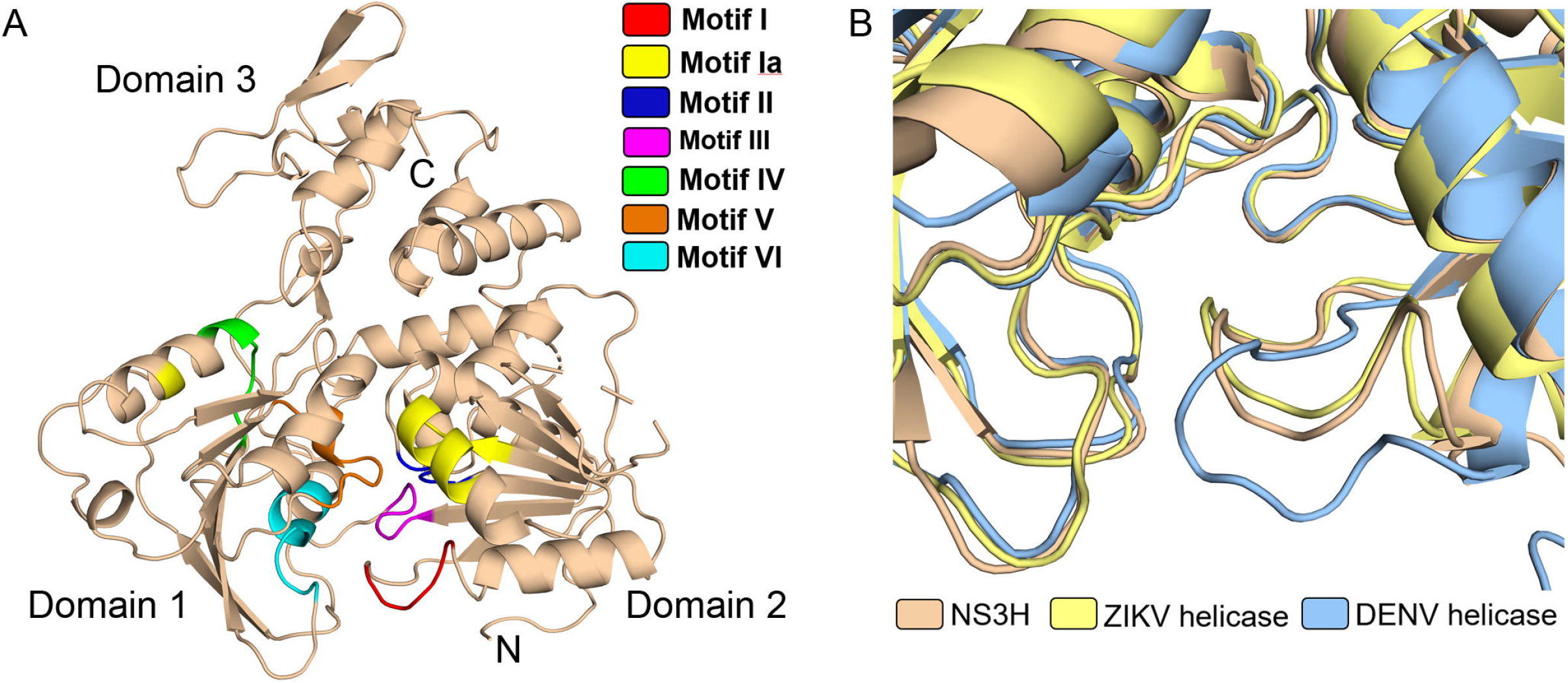
Overall structure of apoNS3H and structural comparison with DENV and ZIKV helicases. (A) A 3-dimensional structure of apoNS3H shown in cartoon representation. SF2 helicase motifs are highlighted in colors. The N- and C-termini are labeled. (B) Superposition of NS3H (wheat) with DENV (light blue) and ZIKV (pale yellow) helicases showing ATPase site. NS3H adapts similar P-loop conformation as ZIKV helicase.

The NTPase site is situated at the interface between domain 1 and 2. It encompasses Walker A and Walker B motifs identified in domain 1, and motif VI in domain 2. Walker A motif consists of G_202_SGKT_206_ sequence forming the P-loop region of NTPase site with residue K205 known to recognize the β- or γ-phosphate of ATP in homologous helicases (12). Walker B motif contains a conserved D_290_EAH_293_ sequence, which is common among flavivirus helicases [reviewed in (4)]. Motif VI (Q_459_RRGRVGR_466_) includes the arginine fingers (R463 and R466), which are important in the energy coupling during the NTP hydrolysis (12,43) and in recognizing the γ-phosphate of ATP (13). The empty NTPase site is filled with solvent molecules and the P-loop adopts a “relaxed” conformation as found in ZIKV structure (10) while DENV helicase exhibits yet more open conformation (11) (Fig. 2B).

### c. Structural snapshots of ATP hydrolysis cycle

Three ternary complexes of NS3H with AMPPNP-Mn^2+^, ADP-Pi-Mn^2+^ or ADP-Mn^2+^, respectively, were captured in crystallographic structures. Based on these structures, NS3H seems to follow a common mechanism of ATP hydrolysis cycle in flaviviruses (11,41). Here, the hydrolysis cycle is represented by four states: (i) a pre-hydrolysis, AMPPNP-Mn^2+^-bound state representing the binding of ATP molecule to the helicase; (ii) a post-hydrolysis intermediate, ADP-Pi-Mn^2+^-bound state, (iii) a product dissociation intermediate with Pi released and ADP-Mn^2+^ are still bound to the protein; and (iv) nucleotide-free (apo) product release state which is ready to bind ATP.

The AMPPNP-Mn^2+^- and ADP-Mn^2+^-bound NS3H structures were obtained via co-crystallization of apoNS3H with corresponding nucleotides while the ADP-Pi-Mn^2+^ complex was obtained as result of ATP hydrolysis during crystallization, i.e., Pi remained trapped within the structure. The nucleotide complexes crystallize in *P*4_1_2_1_2 space group with one molecule in the asymmetric unit similar to the apoNS3H, hence structural changes seen in the ternary complexes are not likely due to differences in crystal contacts. Several localized conformational changes within individual domains were observed (Fig. 3A) when the nucleotide complexes were compared to apoNS3H. Upon nucleotide and divalent ion (Mn^2+^) binding, the P-loop swings toward the bound nucleotide (Fig. 3B) with an inward orientation of K_205_ side chain to coordinate the phosphate group (Fig. 4). In AMPPNP-Mn^2+^ and ADP-Pi-Mn^2+^ complexes, the tip of α7 helix moves away from the relative position in apoNS3H (Fig. 3B) in order to accommodate γ-phosphate. This motion is absent in the ADP-Mn^2+^ structure.

**Figure 3.**
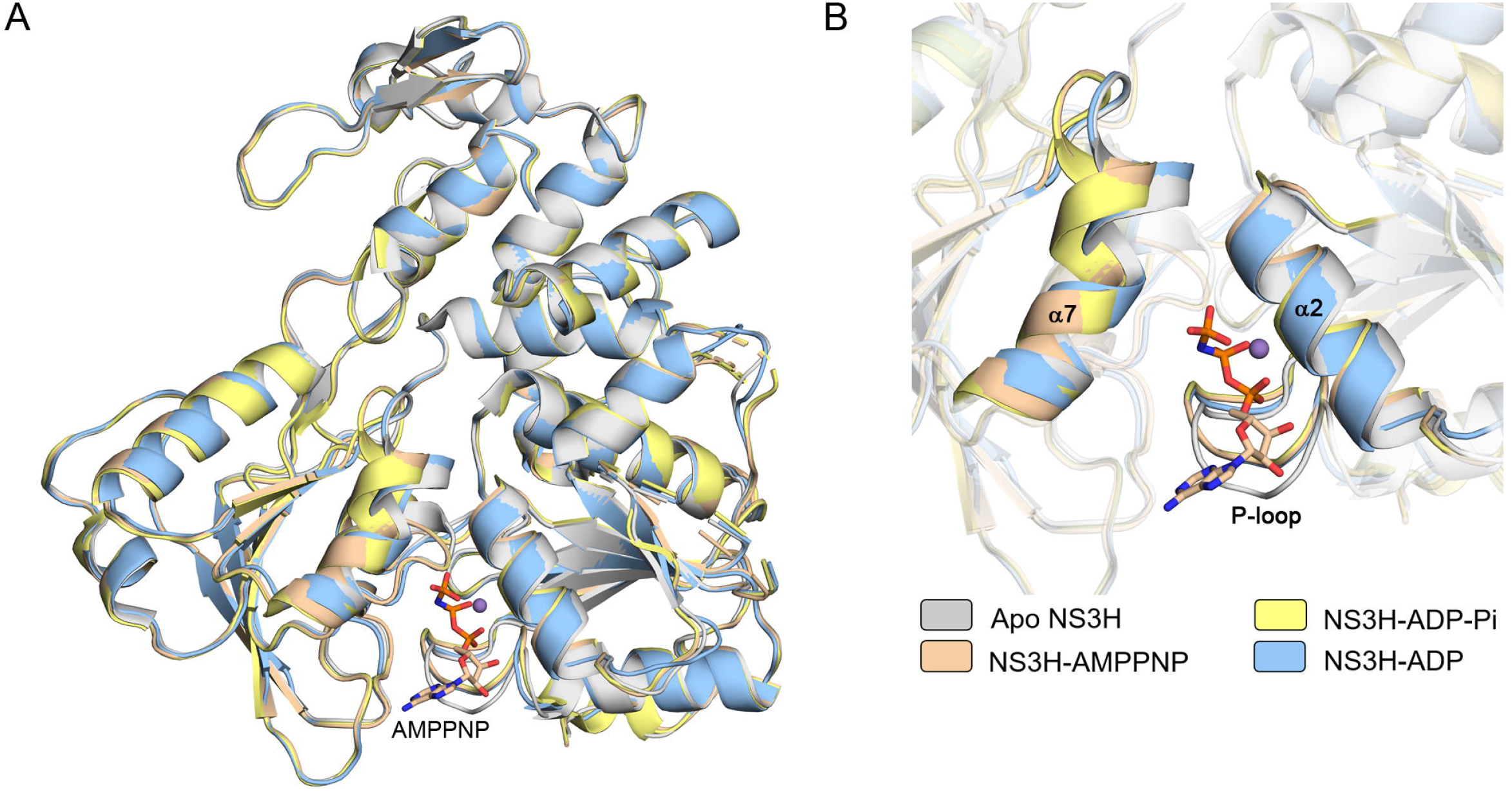
Nucleotide-bound complexes. (A) Superposition of apo and ternary NS3H structures showing global protein conformation. (B) Close-up view of ATPase site corresponding to panel (A) where α2, α7, and P-loop are highlighted. A conformational change in the tip of α7 and P-loop are observed with respect to the bound-ligand. AMPPNP (sticks representation) and manganese ion (purple sphere) are shown.

**Figure 4.**
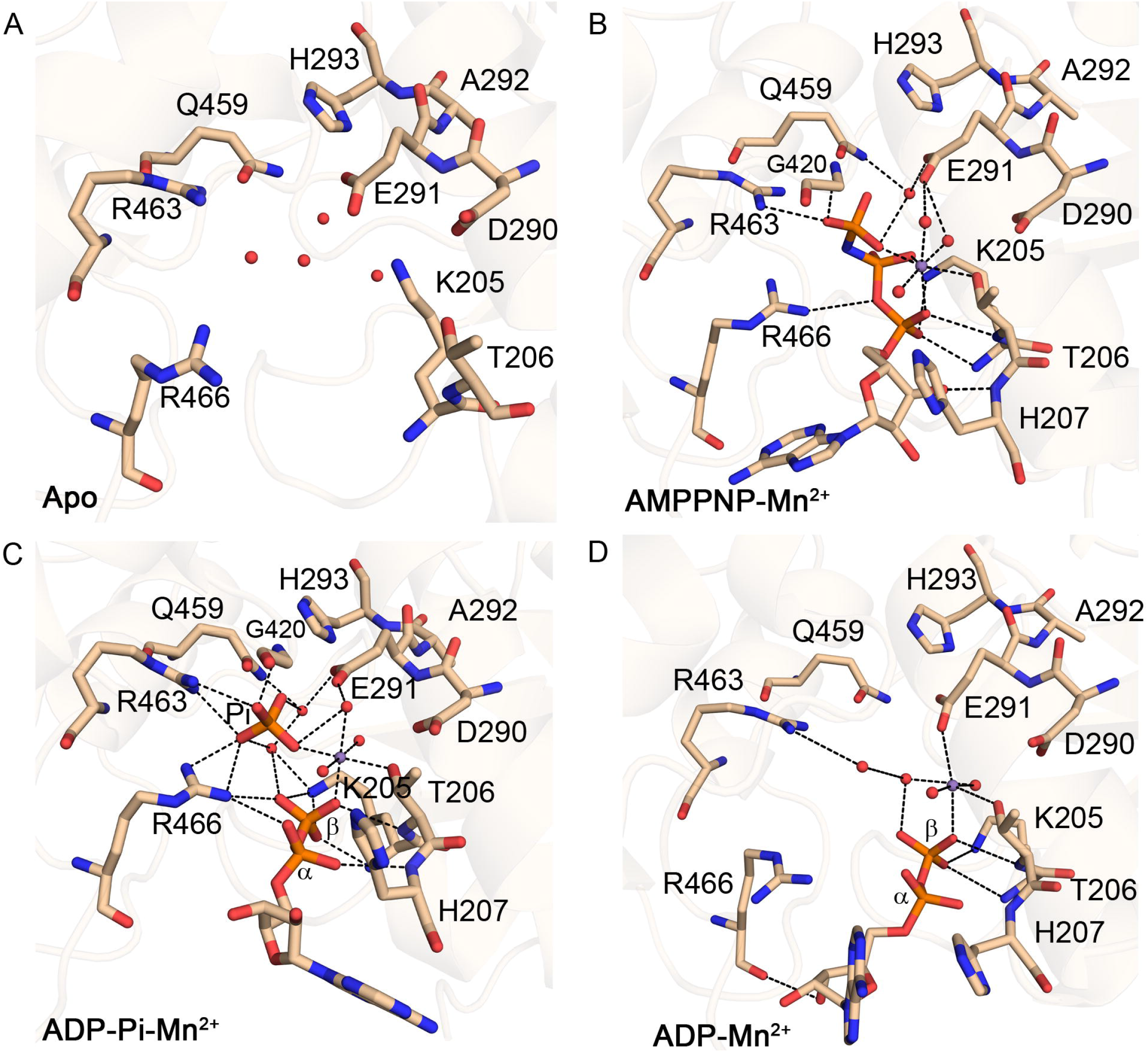
Interactions within nucleotide binding site in apo and nucleotide-bound complexes. Important residues in the NTPase site are shown in sticks representation. (A) apoNS3H, (B) AMPPNP– Mn^2+^ ternary complex, (C) the ADP–Pi–Mn^2+^ ternary complex, and (D) the ADP–Mn^2+^ ternary complex. The manganese ion and water molecules are represented as purple and red spheres, respectively. Hydrogen bonds and metal ion coordination are displayed as black dashed lines.

In the pre-hydrolysis state, NS3H-AMPPNP-Mn^2+^ complex (Fig. 4B), the triphosphate moiety of AMPPNP interacts with P-loop residues K205 and T206 via hydrogen bonds and with residues D290 and E291 of Walker B motif and Q459 from motif VI via water-mediated coordination. One of the arginine fingers, R463, and residue G420 (motif V) also interacts with γ-phosphate of AMPPNP. The 3’-OH group of the ribose C3` endo ring pucker is hydrogen bonded with the main chain amide nitrogen in between residues T206 and H207. One potentially catalytic water molecule coordinates with the side chains of E291 (Walker B), Q459 (motif VI) and γ-phosphorus atom in location suggested by AMPPNP-bound DENV helicase (11) and ATP-bound ZIKV helicase structures (41). The triphosphate moiety of AMPPNP is in a staggered conformation (as defined in 44)) while in DENV helicase (PDB: 2JLR) (11) it adopts a coplanar/eclipsed conformation (Fig. S3), which is proposed to facilitates nucleophilic attack during hydrolysis (44).

The NS3H-ADP-Pi-Mn^2+^ complex (Fig. 4C) was captured following the co-crystallization of NS3H and ATP implying that ATP was hydrolyzed during crystallization or within the crystal and that the subsequent Pi release is extremely slow given the lengthy crystallization protocol. This suggests that phosphate release is the rate-limiting step of the ATPase cycle (in the absence of RNA) and is prevented by the closed conformation of the ATP binding site. The low basal ATPase activity, as measured by phosphate release assay in the absence of RNA, further indicates that this step is facilitated by RNA binding and might be coupled to RNA translocation. This is similar to the proposed HCV mechanism (17). However, recent theoretical results for DENV helicase suggested the ATP hydrolysis as the rate-limiting step, which is accelerated by RNA (9). We observed that both arginine fingers (R463 and R466) and residue G420 (Motif V) directly interact with the free Pi while residues Q459 and E291 interact with the free Pi via water-mediated coordination. It is worth noting that free Pi occupies the same position as γ-phosphate in the AMPPNP-Mn^2+^ complex while the β-phosphate from ADP molecule occupies the position of α-phosphate in AMPPNP-Mn^2+^ complex. On the other hand, the ADP molecule is in similar position as the one captured in ADP-Mn^2+^ complex (Fig. 4D). Thus, the strong interactions of the γ-phosphate are driving nucleoside diphosphate deeper into the binding pocket prior to hydrolysis. Hydrolysis then leads to irreversible phosphate separation, producing a long-lived post-hydrolysis state before Pi and ADP release from the NTPase site.

In the NS3H-ADP-Mn^2+^ complex (Fig. 4D), the arginine fingers do not interact directly with any of the phosphate moiety of ADP. Residue R463 coordinates a water molecule while the side chain of residue R466 moves away from the hydrolysis site. In addition, the 3’-OH group of the ribose C3` endo ring pucker interacts with the main chain carbonyl oxygen of R466. Therefore, this structure captured a state with diminished diphosphate interactions that may facilitate ADP release.

### d. Conformational changes upon RNA binding

Our data and previously published studies (38) show that RNA binds with nanomolar affinity and stimulates NS3H ATPase activity. We followed crystallization strategies similar to those previously used for DENV (11) and ZIKV (40) but with no success. To gain further insight into RNA binding, we constructed a model based on the closely related DENV helicase crystal structure (PDB: 2JLU (6)) with bound (resolved) hexanucleotide ssRNA. RNA oligo was placed into the binding groove by superposing the conserved protein structures. Any clashed in the resulting complex were relieved by molecular mechanics and then subjected to molecular dynamics simulations (trajectories up to 1 μs) to generate ensemble of RNA-bound structures (Fig. 5). RNA interaction energies (E_int_) were computed for frames along the trajectory using generalized Born surface area method (GBSA). Frames with diminished E_int_ may represent states in which weak interactions between the protein and RNA facilitate relative movement and enable translocation. In addition, structural comparison between states exhibiting high and low E_int_, respectively, may reveal how protein conformational changes modulate RNA affinity.

**Figure 5.**
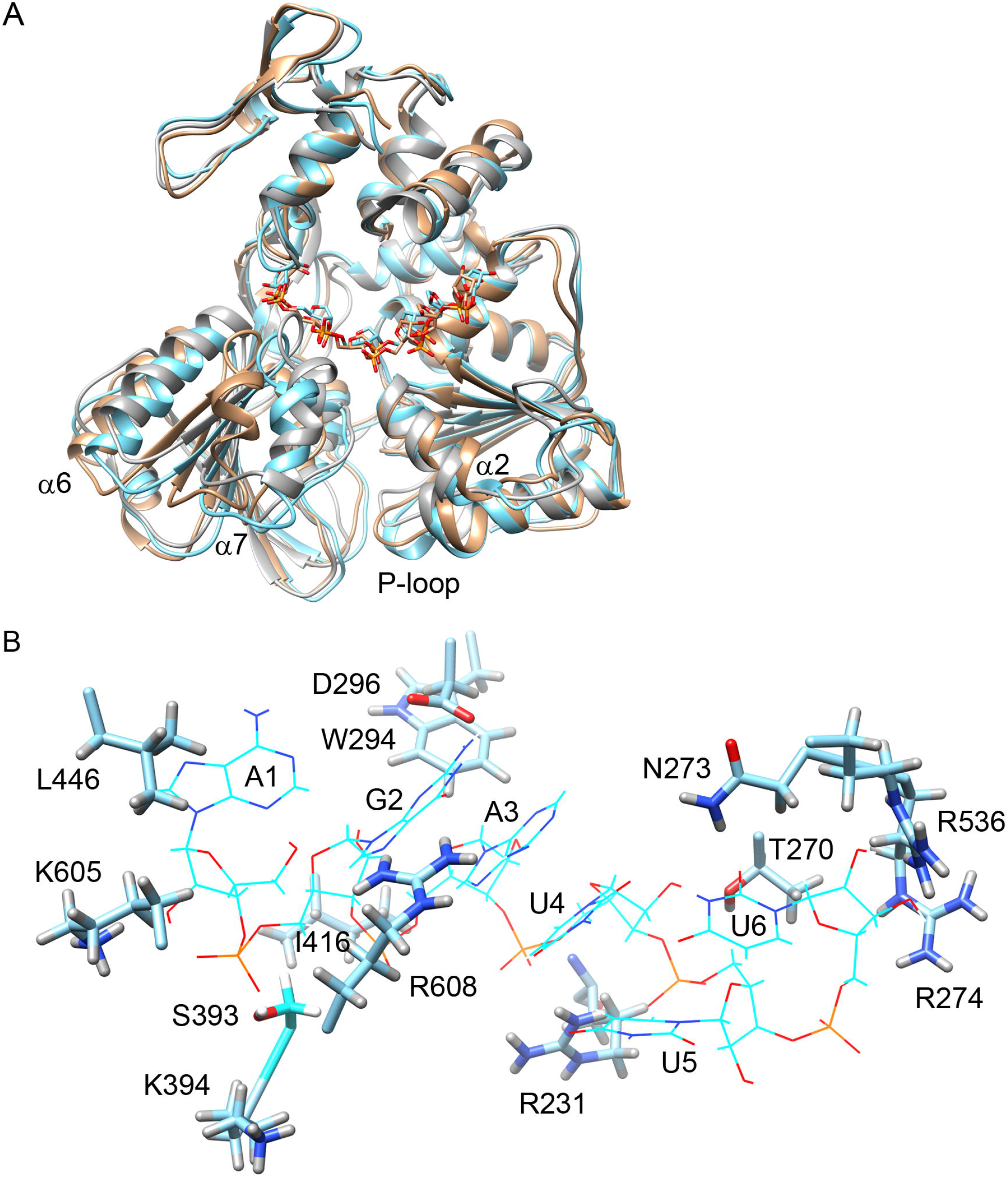
Modeling RNA binding by MD simulations. (A) Comparison of apo crystal structure (grey ribbons) and two RNA-bound conformations with low (wheat) and high (cyan) affinity. Bound RNAs shown as backbone only (two closely related configurations of hexanucleotide). Protein elements that undergo conformational changes associated with RNA binding are labeled in black. (B) Configuration of RNA (wireframe) in the binding cleft of the high-affinity conformation. Interacting protein side chains are shown in stick representation. Carbon atoms-cyan, hydrogen – light grey, oxygen – red, nitrogen – blue, phosphorus-orange.

Both protein and RNA quickly depart from the initial model as demonstrated by RMSD (Fig. S4A, left panel) and sample multiple conformations. Likewise, RNA is bound with widely different E_int_ suggesting substantially dynamic association (Fig. S4A, right panel). Overall structural differences are illustrated by comparing selected frames with low (E_int_ = -150 kJ/mol) and high affinity (E_int_ = -350 kJ/mol), respectively, with the starting apoNS3H structure (Fig. 5A). Overall disposition of protein secondary structure elements and motifs is similar between high RNA affinity and apoNS3H structures while the low affinity conformation differs substantially. With the exemption of the P-loop, which assumes conformation seen in nucleotide containing structures in the presence of RNA, larger conformational changes are mostly confined to domain 2. However, α2 helix in the low affinity structure moves closer to the β-sheet core and opens the nucleotide binding pocket. In domain II, α6 helix shifts away from the RNA binding groove. In the low affinity structure this shift is further augmented by a relative rotation of the whole domain 2. This rotation and further remodeling of α7 helix leads to a widely open nucleotide binding cleft in the low affinity structure while in the high affinity structure α7 helix only rotates around its axis. Slight repositioning of domain 3 further opens the RNA binding cleft in the low affinity structure.

The overall conformational differences are result of local interactions between RNA and protein. As shown in Table 2 the interactions are distributed over residues belonging to all three domains. As expected, there are fewer or weaker interactions in the low affinity structure. In the high affinity model (Fig. 5B), ssRNA backbone is anchored by interactions between polar or charged residues along the tunnel (K605, R231, T270, R536, R274, S393). The polar group interactions are augmented by Van der Waals contact from hydrophobic groups lining the groove (L446, W294, and I416). In addition, there are several base specific contacts. The G2 base, which is conserved within the 5’-UTR of most flaviviruses (15), is recognized by a hydrogen bond between D296 side chain carboxy group and the guanine amino group, and by two hydrogen bonds from the guanidinium group of R608 to N7 and O2 of the G2 base, respectively. Another specific hydrogen bond is between N273 and O2 of U6 base, which is also conserved among many but not all flaviviruses.

**Table 2.** RNA interactions by residue and protein conformation.

| **Protein** | **Atom/group** | **RNA** | **Group** | **Type** | **High affinity** | **Low affinity** |
| --- | --- | --- | --- | --- | --- | --- |
| L446 | CD1 | A1 | base | VdW (Van der Waals) | + | - |
| K605 | NZH | A1 | 2'O | HB | + | - |
| K394 | NZH<sub>3</sub><sup>+</sup> | G2 | P | Salt bridge/ion pair | - | + |
| R608 | NH1/NH2 | G2 | N7/O6 | HB | + | - |
| S393 | OGH | G2 | PO | HB | + | +/- |
| D296 | O | G2 | NH2 | HB (hydrogen bond) | + | + |
| W294 | CH2 | A3 | H4' | VdW | + | + |
| I416 | CD | A3 | 2H5' | VdW | + | +/- |
| T270 | OG | U4 | 2'OH | HB | + | + |
| R231 | backbone NH | U4 | PO | HB | + | + |
| R231 | NH2 | U5 | PO | HB/salt bridge | + | + |
| N273 | NDH | U6 | O2 | HB | + | +/- |
| R536 | NH2 | U6 | 2'O | HB | + | - |
| R274 | NH2 | U6 | P | salt bridge | + | + |

Comparison between the pattern of RNA-protein interactions between the low and high affinity structures offers insights into possible modulation of RNA affinity by protein conformational changes (Fig. S4B and Table 2). In both TBEV models the bound RNA is in an extended conformation (Fig. S4B), similar to that observed in corresponding DENV crystal structure (Fig. S4C). The 5’ end of RNA backbone is anchored in approximately the same place, but the interacting residues are different. In the high affinity structure, the G2 5’ phosphate is held in place by K605 (domain 3) and S393 (N-terminal end of α7 helix). The former interaction is replaced by K394 (N-terminal end of α7 helix) and the S393 contact is weakened in the low affinity structure. The conserved G2 base is shifted in position and only held by D296 and the backbone and base positions downstream are displaced and held by fewer contacts in the low affinity structure (Fig. S4B and Table 2). This suggests that the low affinity structure may represent an intermediate in which domain 2 moves towards the 5’ end while the domain 1 interaction with RNA is weakened and the empty nucleotide binding cleft is wide open. Closure of the nucleotide binding cleft upon ATP binding might then cause movement of domain 1 towards 5’ end, completing an “inchworm” step in the direction of translocation.

Based on our models TBEV NS3H binds RNA in an extended conformation, primarily via interactions with the phosphodiester backbone and in a similar fashion as shown for DENV and ZIKV helicases (11,15,40). RNA specificity is augmented by interactions of domain 1 T270 with 2’OH of U4 and domain 3 R536 hydrogen bond to 2’O of U6 (Table 2). Furthermore, specific recognition of the 5’-UTR sequence is primarily mediated by three hydrogen bonds (D296, R608) to G2 and augmented by N273 hydrogen bond to O2 of U6 base, which is also part of the conserved 5’-UTR motif. Apart from D296 the sequence specific interactions are weakened in the low affinity conformation, and this is similar to the alternative conformations of R538 seen in crystal structures DENV helicase with different RNAs (15). Unlike crystal structures, in which larger domain motions might be prevented by lattice packing, MD simulations suggest that the protein is quite dynamic in the RNA-bound complex and explores conformations with different RNA affinity and disposition of domain 1 and 2. These configurations may facilitate handover of RNA between these two domains as proposed by the “inchworm” mechanism (45) and “rachet mechanism” (46). It remains to be seen whether and how are these configurations related to phosphate release, the rate-limiting step of the ATPase.

### e. Structural basis of DNA inhibition

Biochemical data show that ssDNA acts as an inhibitor of RNA-stimulated ATPase activity. The inhibition is incomplete (Fig. 1D) and requires much higher ssDNA concentrations (micromolar range) than suggested by RNA affinity (*K*_*D*_ ∼ 50 nM, Fig. 1E), thus suggesting DNA does not directly compete with RNA for the binding cleft. This is further supported by only partial binding competition even at high DNA concentrations (anisotropy, Fig. 1F).

Furthermore, the increased effectiveness of longer ssDNA_41_ suggests that binding to multiple surface sites might be involved. The electrostatic surface charge distribution (NS3H:RNA model, Fig. S5) suggests that even in the presence of RNA there are positively charged patches around the ATP binding site and near the exit site of the RNA groove on domain 1. Hence, DNA binding along the ATP binding cleft might be a plausible way to interfere with the ATPase cycle, e.g., by biasing the conformation of the ATPase site toward inactive configuration or modulating the ATP affinity.

We have explored this hypothesis by constructing a model based on the NS3H:AMPPNP structure (with AMPPNP replaced by ATP) and adding a short (hexanucleotide) DNA with B-form backbone running along the ATP binding cleft. The initial model was subjected to MD and rapidly converged to a stable ensemble of configurations (Fig. 6A). As controls two MD simulations were done: one without the bound DNA and another starting from the last frame of the NS3H:ATP:DNA complex but with the bound DNA removed. Analysis of ATP interaction energy distributions obtained from the stable sections of these stimulation trajectories (as judged by RMSD, Fig. S6A) revealed that DNA binding along the ATPase active site reversibly decreases the apparent ATP affinity by about two-fold. Similar effect is observed when ssDNA_41_ is placed across the cleft (Fig. 6C). In this case the DNA, which in the initial model is represented by a long, straight B-type helix, folds and bends and interacts with multiple positively charged patches (*c.f*. surface charge map in Fig. S5). While binding along the ATPase cleft seems persistent, the interactions with positive patches II and III are transient on the nanosecond time scale (Movie S1).

**Figure 6.**
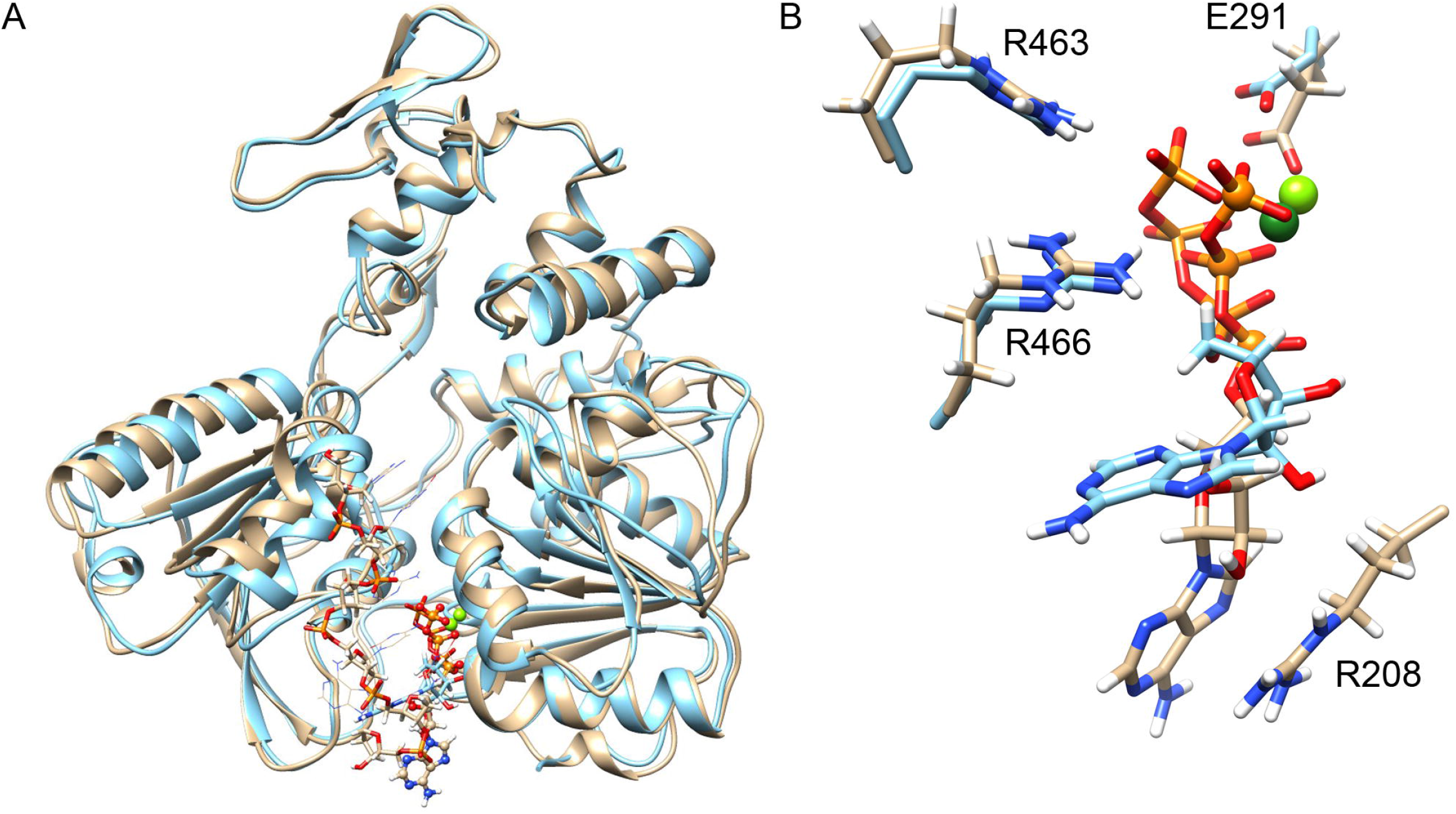
Effects of short ssDNA association on ATP binding. (A) The conformational changes upon DNA binding (final conformation, bases in wireframe and backbone in sticks representation), ATP is shown in sticks (initial, cyan) and ball-and-stick (final, wheat) representation. Protein backbone is shown as a ribbon (final state wheat). Green sphere indicates position of Mg. (B) Close-up of the ATPase active site showing γ-phosphate displacement. (C) Change in ATP affinity upon DNA binding measured by violin plot of GBSA interaction energy (-E_int_) distributions. Short DNA refers to hexanucleotide sequence 5’-AGACTA-3’.

Given that ATP binding is dominated by polar and electrostatic interactions with the triphosphate moiety and the adenine base is highly mobile (as seen in MD simulations) it is not surprising that the DNA-induced decrease in interaction energy is associated with displacement of the triphosphate moiety within the ATPase active site (Fig. 6B and Fig. S6B). Such displacement would be enough to disrupt the precise alignment of the catalytic residues, water and the γ-phosphate thus may lead to inhibition of the ATPase cycle. DNA binding may also interfere with nucleotide exchange by sterically blocking access to the ATPase site and the longer DNA also partially occludes the RNA binding groove.

## 4. Conclusion

Current study provides a structural and mechanistic insight into the ATP hydrolysis cycle of TBEV helicase. Several conformational changes associated with individual ATP hydrolysis steps have been captured in crystallographic structures of NS3H-AMPPNP-Mn^2+^, -ADP-Pi-Mn^2+^, -ADP-Mn^2+^ and the apo form of the protein. Overall structure of TBEV helicase is highly similar to that of other flaviviruses such as DENV and ZIKV, suggesting similar ATP hydrolysis and RNA translocation mechanism. However, contrary to mechanism proposed for DENV helicase (9), our structural and biochemical data is consistent with Pi release being the rate-limiting step of the ATPase, as seen also for the HCV NS3 helicase (17).

Furthermore, we found that ssDNA inhibits RNA-stimulated ATPase activity but not by direct competition with RNA for the oligonucleotide binding cleft. Using modeling and MD simulations we propose a plausible explanation in which ssDNA exploits nonspecific binding to positively charged surface patches in the vicinity of the ATP binding pocket, weakening ATP interactions, and consequently leading to repositioning of the triphosphate moiety. However, this is just one plausible explanation and binding of longer DNAs might have additional effects on protein dynamics, e.g., preventing conformational changes that are necessary for phosphate release or nucleotide exchange. While ssDNA is irrelevant to the virus replication cycle in the cytoplasm where it is unlikely to encounter this type of nucleic acid, it might open new ways to design polyanionic allosteric inhibitors that combine this “parasitic” mode with more specific targeting. These remains to be elucidated in the future studies.

## Acknowledgments

We are grateful to the help and support of Drs. Pavlína M. Řezáčová, Jiří Brynda, and Manfred Weiss during diffraction data collection in BESSY II electron storage ring operated by the Helmholtz-Zentrum Berlin. We would also like to acknowledge Dr. Filip Dyčka for his contribution in MALDI-TOF mass spectrometry, and Petra Havlíčková and Dr. Pavel Grinkevich for helpful discussions and contribution during structural refinement.

## Funding and additional information

This work was supported by ERDF CZ.02.1.01/0.0/0.0/15_003/0000441.

