## supplemental figures for "Mechanistic insight into the ATP hydrolysis cycle of tick-borne encephalitis virus helicase"

**Contents:**

**Table S1.** List of oligonucleotides used in wet experiments
**Figure S1.** Purification of recombinant NS3H.
**Figure S2.** Structural-based sequence alignment of NS3 helicases.
**Figure S3.** Comparison between α7 and AMPPNP molecule position from AMPPNP-$Mn^{2+}$-bound NS3H and DENV helicase (PDB: 2JLR).
**Figure S4.** MD simulation.
**Figure S5.** Surface electrostatic energy distribution for NS3H:RNA complex model.
**Figure S6.** MD simulation involving NS3H and DNA.

**Table S1. List of oligonucleotides used in the experiments**

| | |
|---|---|
| **Primer for cloning into pET-19b vector** | |
| NS3Hel_NdeI_F[a] | 5'-TGCTAGCATATGGAGAAGAGTCGACCCAACCTCCC-3' |
| NS3Hel_XhoI_R[a] | 5'-TGCTAGCTCGAGTTAGCGACGCCCAGATGCGTA-3' |
| **Labeled ssRNA for fluorescence anisotropy binding assay** | |
| 6-FAM-ssRNA$_{12}$ | [6-FAM]-5'-AGAUUUUCUUGC-3' |
| **Unlabeled ssDNA for ATPase/fluorescence anisotropy binding assays/MD simulation** | |
| ssDNA$_{6}$ | 5'-AGACTA-3’ |
| ssDNA$_{12}$ | 5'-AGATTTTCTTGC-3' |
| ssDNA$_{20}$ | 5'-AGAGGATCCCCGGGTACCGA-3' |
| ssDNA$_{41}$ | 5'-AGAGGATCCCCGGGTACCGATTAGATTATTGAGCTCTCCAG-3' |

[a] Restriction enzyme sequence is underlined.

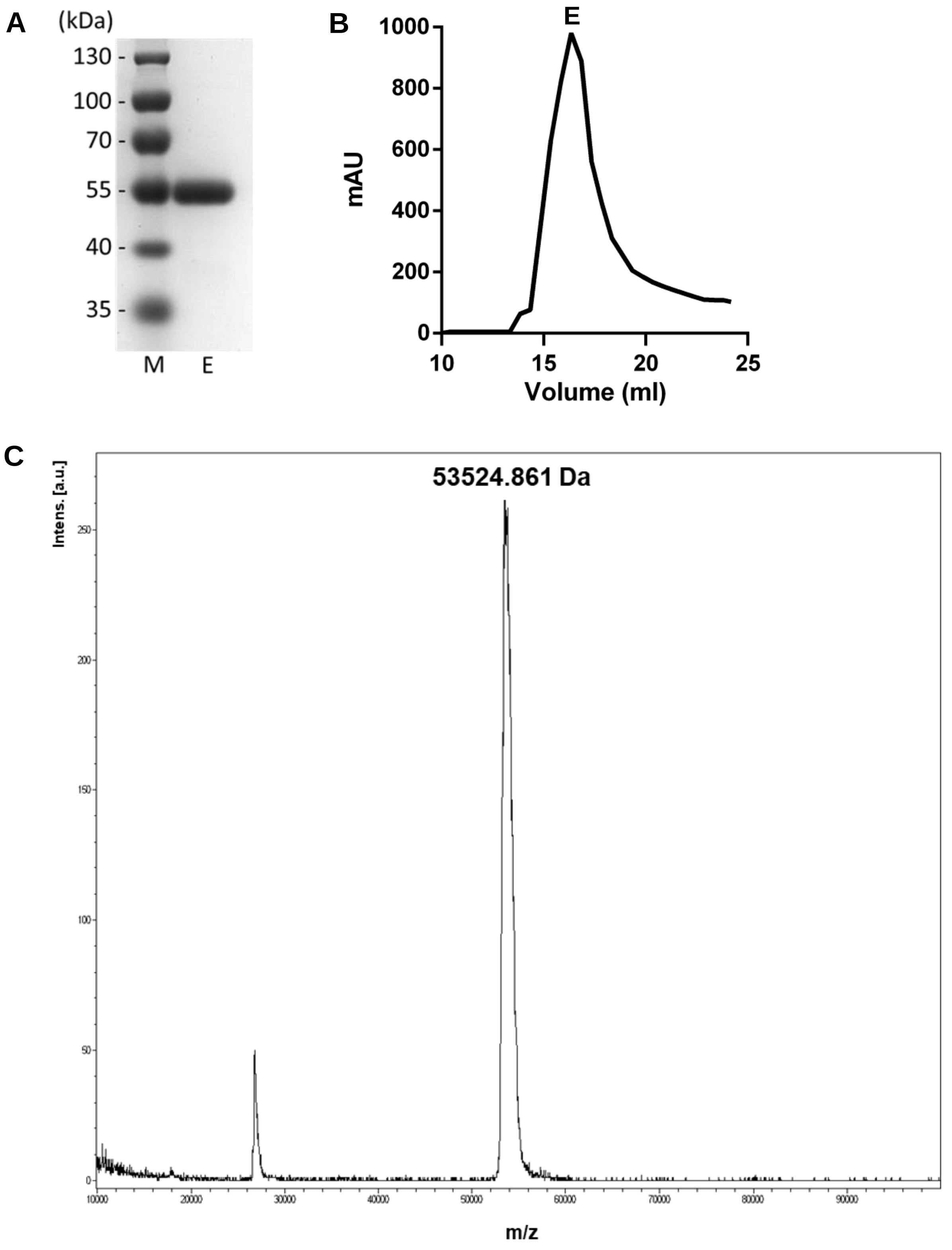


**Figure S1. Purification of recombinant NS3H.** (A) An SDS-PAGE analysis of eluted recombinant protein after gel filtration chromatography. Lane M is the SDS-PAGE molecular weight standard and lane E indicates the elution peak from gel filtration chromatography. (B) Chromatogram from gel filtration. Letter E indicates the elution peak correspond to panel (A). (C) MALDI-TOF spectrum of purified monomeric NS3H (~53 kDa), confirming the purity and size of recombinant NS3H. Note that the peak at m/z ~27,5 kDa represents a +2 species.

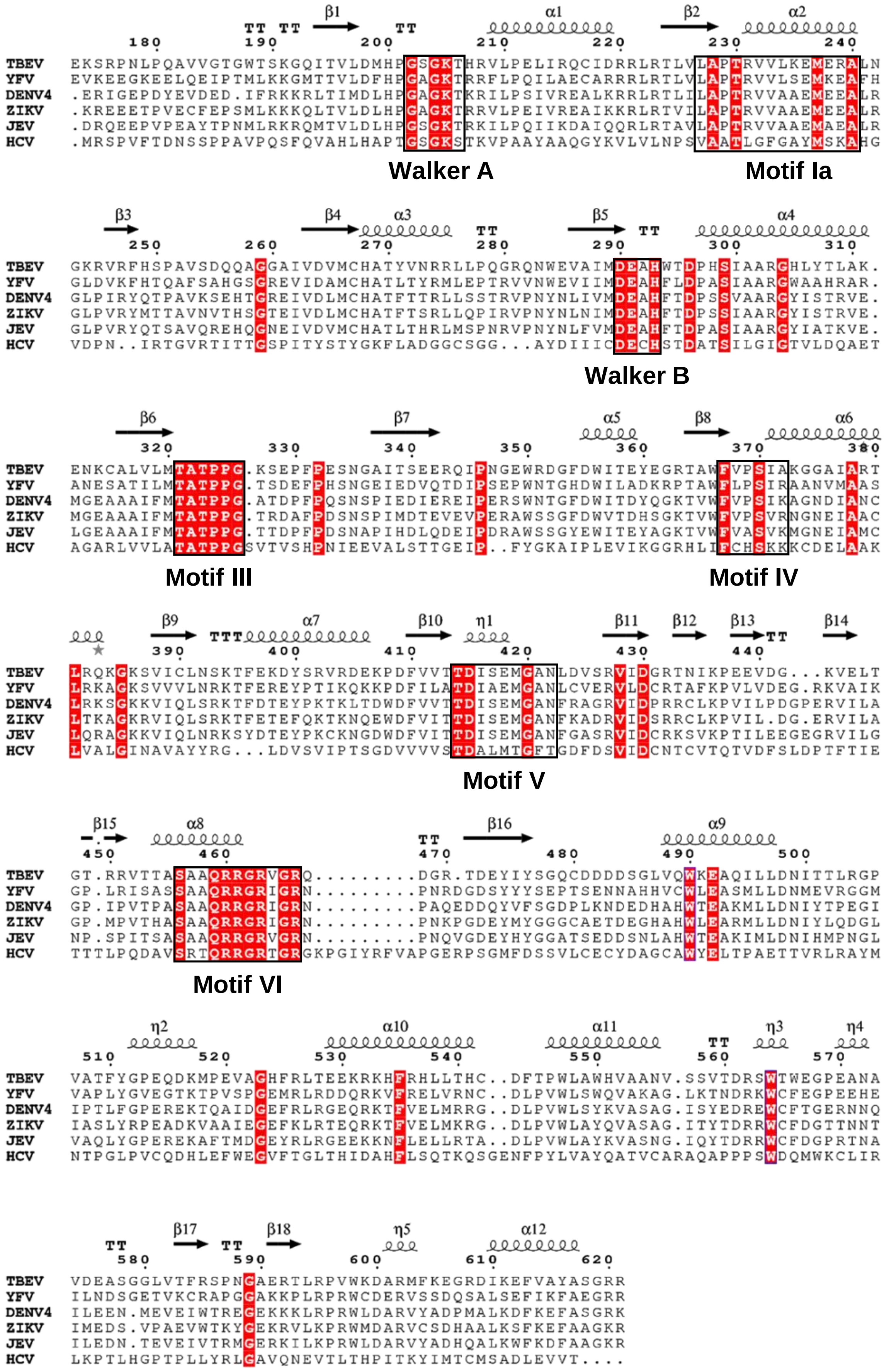


**Figure S2. Structural-based sequence alignment of NS3 helicases.** The alignment was generated using ESPript 3.0 (https://espript.ibcp.fr) for flaviviruses (TBEV, YFV, DENV4, ZIKV and JEV) and HCV. Conserved motifs are labeled and highlighted in a black box.

Reference: Robert, X. and Gouet, P. (2014) "Deciphering key features in protein structures with the new ENDscript server". *Nucl. Acids Res.* 42(W1), W320-W324.

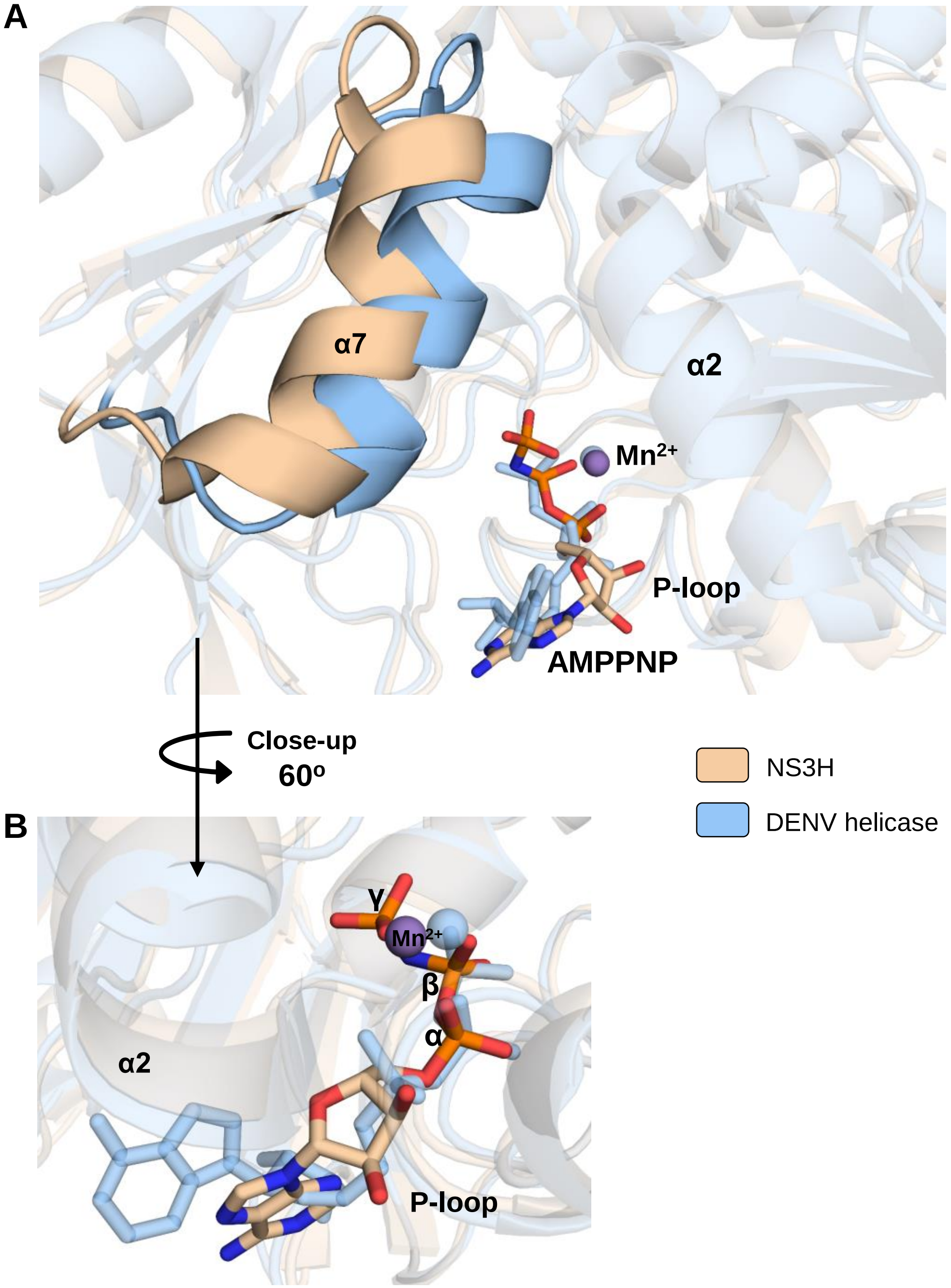


**Figure S3. Comparison between α7 and AMPPNP molecule position from AMPPNP-$Mn^{2+}$-bound NS3H and DENV helicase (PDB: 2JLR).** (A) Superposition of α2, α7, and P-loop conformations from NS3H and DENV helicase. (B) Close-up view of ATPase site highlighting different position of AMPPNP molecule in NS3H and DENV helicase. Manganese ions are shown as purple (NS3H) and blue (DENV helicase) sphere.

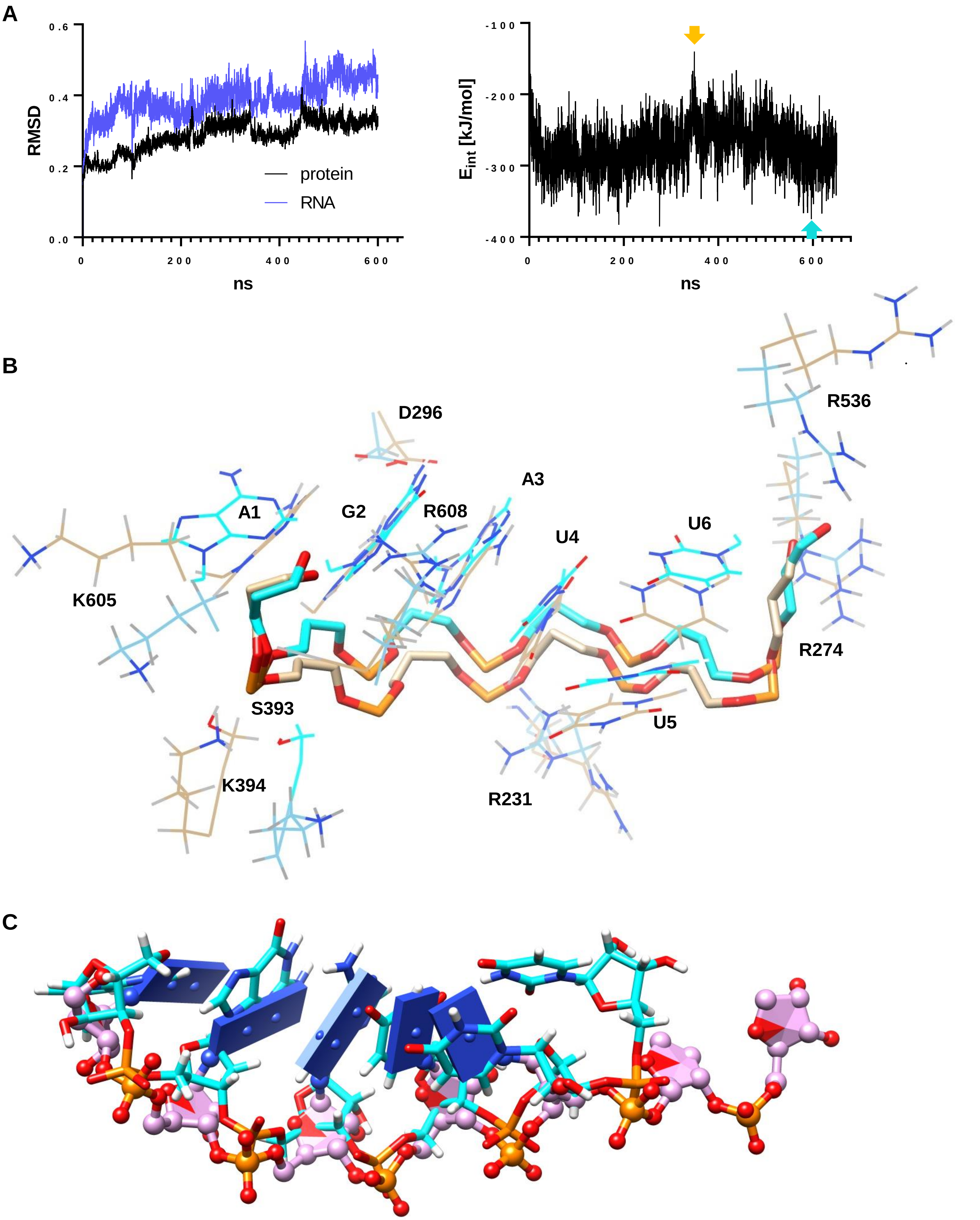


**Figure S4. MD simulation.** (A) Frame RMSD (left) and corresponding RNA interaction energy (GBSA approximation, right). (B) High (cyan) and low (wheat) affinity RNA-protein interaction comparison. RNA backbone in sticks, bases and protein side chains in wireframe. (C) Comparison between RNA configuration in DENV (PDB: 2JLU, in magenta and ball-and-stick representation) and high (cyan, sticks) affinity RNA configuration.

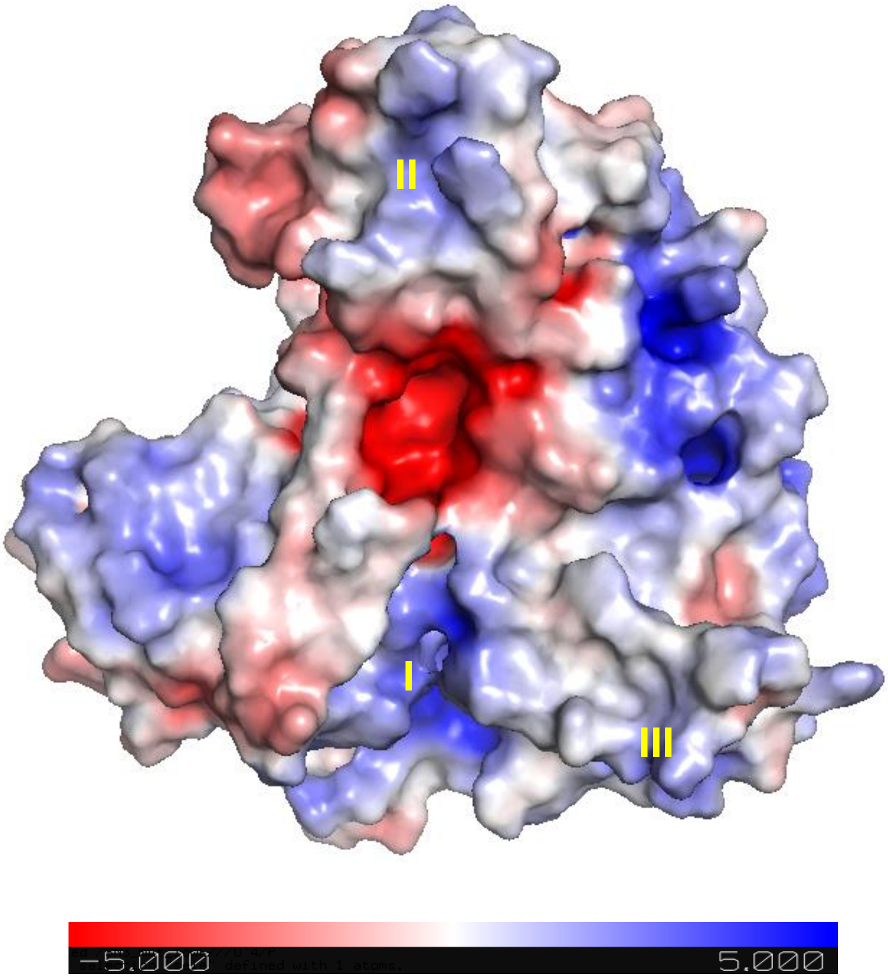


**Figure S5. Surface electrostatic energy distribution for NS3H:RNA complex model.** Positively charged surface areas that might bind additional nucleic acid are numbered (I-III) in yellow.

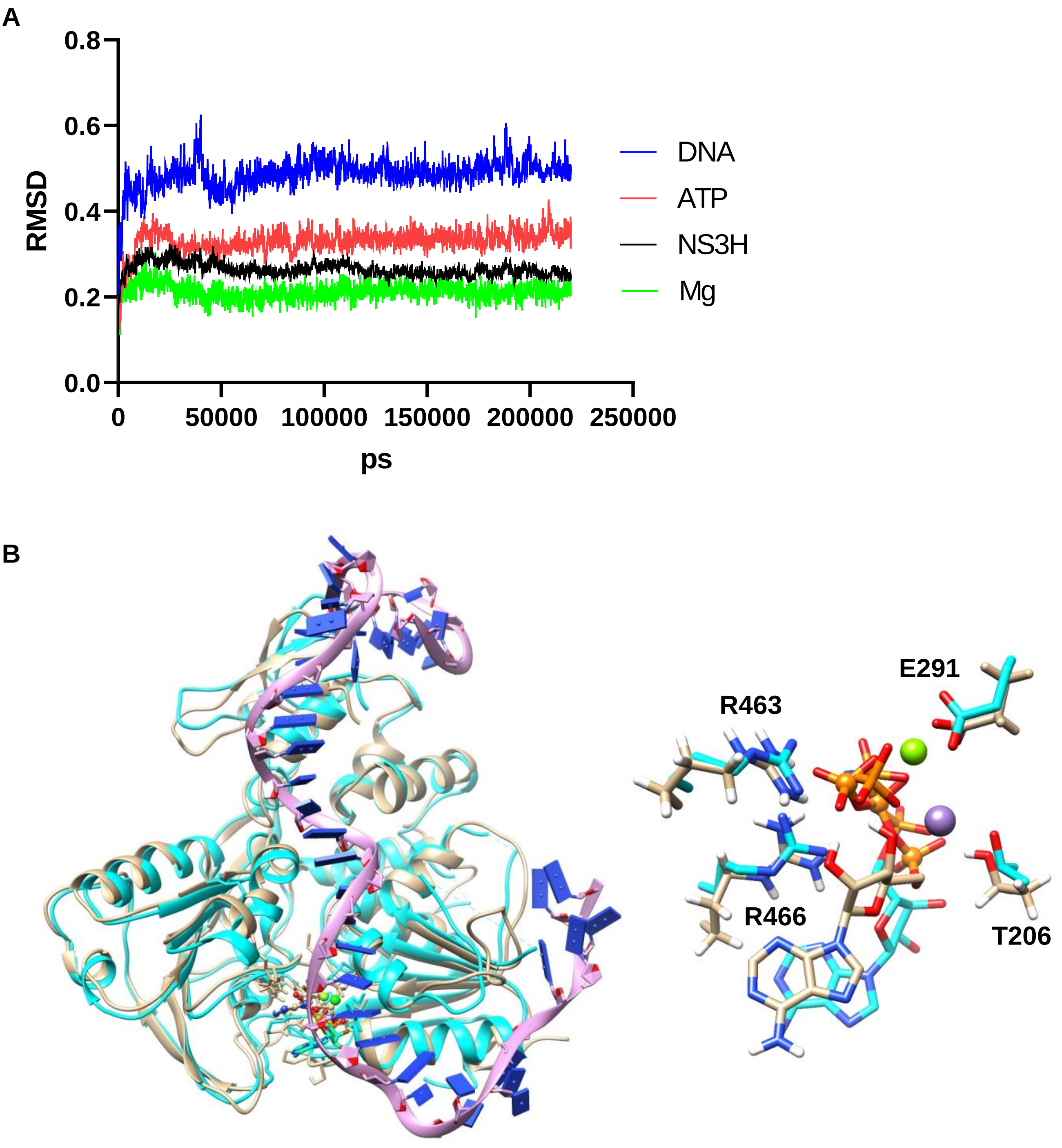


**Figure S6. MD simulation involving NS3H and DNA.** (A) RMSD of NS3H:ATP:Mg complex with short (hexamer) DNA bound along the ATPase site. (B) Left panel: A representative frame from simulation of $ssDNA_{41}$ (magenta backbone, blue bases) associated with NS3H:ATP:Mg complex. The initial state, NS3H:AMP-PNP:Mn crystal structure, is depicted in cyan while the final simulation frame is depicted in wheat. Right panel: Detain of ATP binding upon $ssDNA_{41}$ binding (initial state is cyan, final state is wheat, triphosphate in final state is depicted as ball & sticks). Mn atom is depicted as purple sphere and was replaced by Mg for simulation (green sphere, final state).
